# Rough Grouping Enhances YOLO’s Pollinator Classification and Detection from Small Datasets

**DOI:** 10.1101/2024.08.24.609510

**Authors:** Suyeon Kim

**Author notes:** Corresponding author: Suyeon Kim.

## Abstract

This study addresses the challenges in pollinator monitoring by proposing an effective data structure for automated systems, with a focus on the use of machine learning to handle underrepresented groups in small datasets. By experimenting with grouping the top three pollinators (bee, butterfly, hoverfly) and non-pollinators in datasets of fewer than 300 samples, the research aims to enhance classification and detection accuracy. During 4-hour filming sessions, 181 images of insects larger than 1 cm were captured and classified into three grouping methods: “Pollinator/ Non-pollinator”, “Bee/Butterfly/Hoverfly/Ant”, and “Bumblebee/Honeybee/Butterfly/Hoverfly/ Ant”. YOLO V8 models were trained, validated, and tested with these datasets based on different class grouping methods. The study found that the “Pollinator/Non-pollinator” YOLOv8 model performed best across all metrics, suggesting it is more reliable for categorizing groups and detecting target objects, especially with smaller, imbalanced datasets. This finding aligns with the trend in machine learning that providing more training opportunities for individual classes improves accuracy. Therefore, using broader categorization methods can enhance the reliability and accuracy of automated monitoring systems when training data is insufficient.

## 1. Introduction

Pollinators provide essential and irreplaceable ecosystem services that are crucial for humanity (Ameixa *et al*., 2018). They play a vital role in maintaining biodiversity and ecosystem stability, in addition to enhancing crop productivity, farm income, and access to nutritious food (Potts *et al*., 2016; Garibaldi *et al*., 2020).

However, pollinators are facing numerous survival threats, including environmental pollution, climate change, invasive species, habitat loss, pathogens, and pesticides (Cardoso *et al*., 2020; Harvey *et al*., 2022). Despite the severity of pollinator decline, which could lead to an extinction cascade, these crucial guardians of food security remain under-investigated (Huang *et al*., 2021).

This study aims to propose an effective data structure to support an automated pollinator monitoring system. Data collection is critical for investigating pollinators; however, traditional methods, such as pan-trapping, have recently been shown to be ineffective (Howard *et al*., 2021; Pegoraro *et al*., 2020). As an alternative, machine learning models that utilize integrated computer vision and automated species recognition are gaining attention. With the increasing accessibility of this technology, networks of camera monitoring systems could be installed on a large scale. This would enhance our understanding of the factors behind pollinator declines, help develop effective solutions to address this global issue, and provide a comprehensive understanding of pollination dynamics (Howard *et al*., 2021; Pegoraro *et al*., 2020).

Traditional data collection on interactions between organisms and their environment has relied on on-site human observations, which are time-consuming and limit around-the-clock monitoring of wild animals (Steen, 2016). Video monitoring, however, allows for repeated viewing of flower visitors in real-time and slow motion, aiding in accurate species identification, abundance counting, and behavior sampling (Steen and Aase, 2011; Steen, 2012; Steen and Mundal, 2013; Gimenes *et al*., 1996; Helfrich-förster *et al*., 1998; Bellusci and Marques, 2001; Bloch *et al*., 2013). Furthermore, automated identification monitoring not only enables the observation of a spatio-temporally wider range at lower costs, but also allows for the collection of unique data, such as previously unnoticed behaviors, long-term patterns, full species list in a field, and data unaffected by visit times or weather conditions (Pollinator monitoring more than pays for itself ; Laverty, 1980; Manetas and Petropoulou, 2000; Micheneau et al., 2008).

Previous research on pollinator auto-classification or detection monitoring systems has focused on classifying a few species using various AI algorithms. For instance, a study by Barlow *et al*. (2017) manually collected data by observing bees in time-compressed, time-stamped videos filmed by the RANA digital monitoring system to distinguish between pollinator bumblebees and robbery bumblebees. More recent research in 2023 developed an insect tracking and classifying model with a dataset of over 2000 insect tracks from four insect classes, achieving a precision of 0.995, a recall of 0.767, and an F-score of 0.85 (Ratnayake, 2023). Bjerge *et al*. (2023) distinguished nine dominant insect pollinators, achieving an 80% detection rate and classifying them with an average precision of 92.7%, a recall of 93.8%, an mPA.5 of 94.4%, and an mPA.5-95 of 59.2% using YOLO V3/5 models trained on a manually acquired dataset of 2,347,944 images from 10 cameras.

Recently, many studies have focused on methods to enhance model performance, often including preprocessing or post-processing steps. Another study by Bjerge *et al*. implemented a two-step deep learning model using preprocessed RGB images and the Faster R-CNN for object detection, improving the average micro F1-score from 0.49 to 0.71 with the YOLO detector and from 0.32 to 0.56 with the Faster R-CNN detector (Bjerge *et al*., 2023).

However, leveraging grouping strategies to improve machine accuracy with small datasets has not been attempted until now. This is because insect datasets often have biased or underrepresented data volumes for certain species or groups. This bias is even more pronounced in field data, where the natural frequency of different groups can vary greatly. Generally, the performance of machine learning models decreases when groups with fewer training opportunities are included (Shaikhina *et al*., 2015; Zhang and Ling, 2018; Vabalas *et al*., 2019; Kokol *et al*., 2022; Dou *et al*., 2023).Therefore, understanding how to handle underrepresented groups to achieve reliable automated detection and classification is crucial.

In this study, we experimented with grouping the top three pollinators (bee, butterfly, hoverfly) that appear with different frequencies in datasets of fewer than 300 samples. Our aim was to determine the optimal level of grouping that enhances classification and detection accuracy.

## 2. Methods and Materials

### CAMERA DEVELOPMENT

This study utilized two units of Raspberry Pi camera setups, each comprising one HQ Camera (Arducam 8-50mm C-Mount Zoom Lens), HQ module (IMX477), Raspberry Pi 4 Model B, Romoss 30W 20000mAh power bank, and a tripod. A portable monitor was connected to the Raspberry Pi 4 Model B through a micro HDMI to Type B cable when needed.

### DATA COLLECTION

To acquire images of specimens, each HQ camera was focused closely on a couple of flowers to take videos of insects visiting them (Figure 1). The HQ camera was activated through ‘libcamera’ commands on a Raspberry Pi 4 terminal. Filming flowers and their visitors at several different sites included Cornell Botanic Gardens (42.450254661757754, -76.47267753578093) and solar field sites (42.938457, -74.374467; 42.383867, -76.600515).

**Figure 1.**
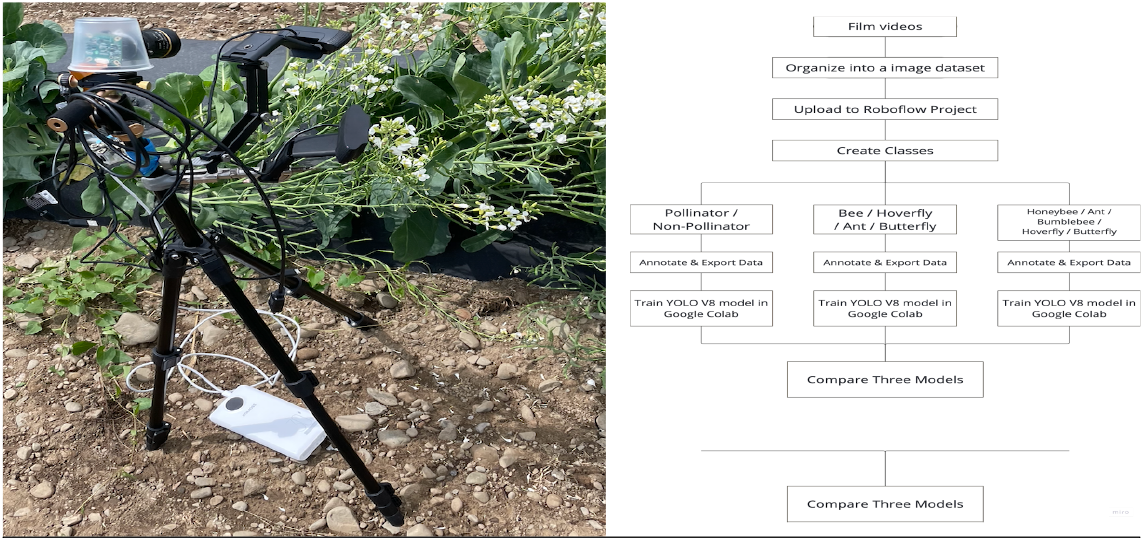
A camera filming a video in the field (Left). The research process after filming (Right).

### CLASS IDENTIFICATION

The insects larger than 1 cm that appeared during the 4-hour filming are as follows: 2 ants, 3 bumblebees, 1 butterfly, 1 fly, 3 honeybees, and 3 hoverflies. On average, 4 different angles of images were acquired from each insect, producing 181 images. When labeling the dataset, three grouping methods were applied: pollinator and non-pollinator; bee, butterfly, hoverfly, and ant; bumblebee, honeybee, butterfly, hoverfly, and ant. For the pollinator and non-pollinator grouping, the experiments were conducted in two ways: one with additional ladybug images downloaded from the internet to compensate for the lack of non-pollinator data, and one without these additional images to maintain the same dataset as other grouping methods for easier comparison. 56 images of ladybugs (*Harmonia axyridis*) were exported from a citizen science project (https://www.lostladybug.org/) to balance the dataset for the comparison of pollinators and non-pollinators grouping. The ladybug images were chosen to have a similar composition and background to the other insects’ images we took ourselves.

The dataset contains 156 images of pollinators and 94 images of non-pollinators. Among the pollinators, bees (102 images), hoverflies (33 images), and butterflies (21 images) were the most frequently captured. In the non-pollinator category, 21 images were classified as ants, and 7 as flies, while the remaining 56 images were classified as ladybugs sourced from the web. Although flies were included in the non-pollinator dataset, they were not used as a separate category due to their insufficient sample size. Ladybug images were solely used for distinguishing between pollinators and non-pollinators.

**Figure 2.**
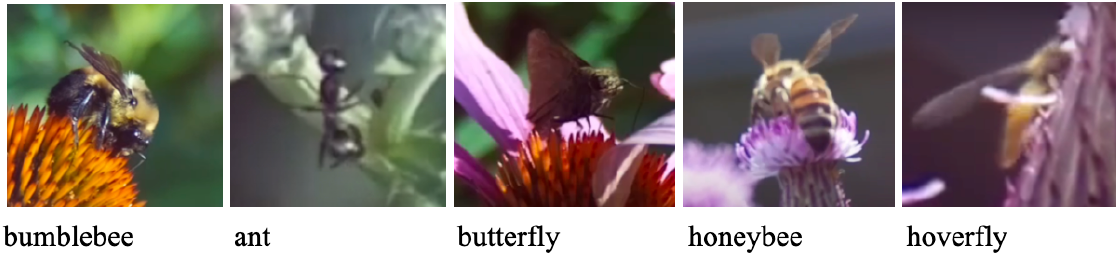
Captured objects categorized by their taxonomic groups from camera recordings.

**Figure 3.**
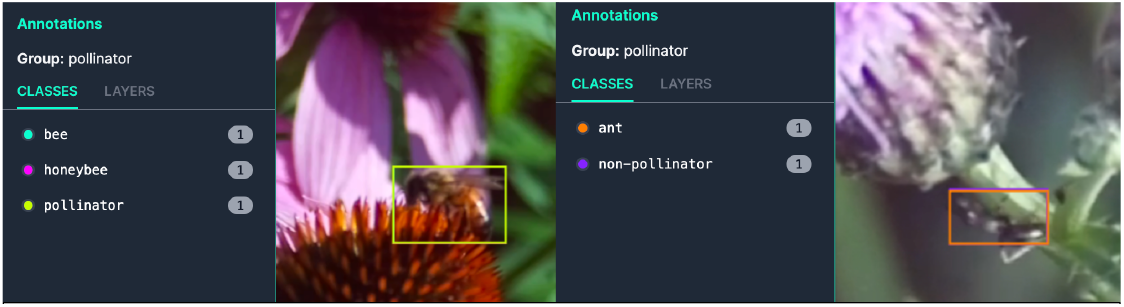
Example of Annotation: The honeybee falls into the “pollinator,” “bee,” and “honeybee” categories simultaneously.

### OBJECT ANNOTATION

To transform the videos into a dataset of images, flower-visiting insects were manually screen-captured. For annotation, the bounding box tool from the Roboflow Object Detection Project was used. The steps of annotation included manually labeling 100 images to activate the auto-labeling function and manually approving or adjusting the automatically labeled boxes.

Classes created for the annotation were “pollinator,” “non-pollinator,” “bee,” “butterfly,” “hoverfly,” “fly,” “ant,” “ladybug,” “honeybee,” and “bumblebee.” Since the pollinator/non-pollinator grouping method is extensive compared to other methods, the “pollinator” class box was always overlaid with the “bee,” “butterfly,” or “hoverfly” box. Similarly, “non-pollinator” was always overlaid with the “ant,” “fly” and “ladybug” class. The “bee” class is comprehensive as well, so the “bee” box was always overlaid with either the “honeybee” or “bumblebee” class.

### MODEL DEVELOPMENT

YOLO V8 models (shared in the Appendix) were trained, validated, and tested with three different datasets based on different class grouping methods In the Google Colab environment, utilizing the functions provided by the ultralytics library. The model trained with a pollinator-or-non-pollinator dataset was expected to detect insects of several different species and classify them into two groups: pollinators or non-pollinators. The objective of the other two models was to detect insects and assign them to more precise taxonomic group names: bee, butterfly, hoverfly, ant, ladybug; bumblebee, honeybee, butterfly, hoverfly, ant, ladybug. The train, validation, and test split ratios were 70:15:15, and accuracy was evaluated using precision, recall, mAP50, and mAP50-95.

## 3. Results and Discussion

### 3.1 Classification of “Pollinator” versus “Non-pollinator”

The YOLOv8 algorithm, trained with pollinator (bumblebee, honeybee, hoverfly, butterfly) and non-pollinator (ant, fly) classes, without web-sourced ladybug images to balance class volume, showed 0.941 in Precision, 0.832 in Recall, 0.880 in mAP50, and 0.490 in mAP50-95 on the test dataset. For both pollinators and non-pollinators, the detection of insect occurrence (mAP50) and classification (Precision and Recall) were relatively high, generally exceeding 80%. This indicates that this model is more reliable in detecting the presence of insects and identifying whether they are pollinators, rather than in the precise detection of their size and location (mAP50-95).

**Table 1.** Validation and Test Performance of “Pollinator” versus “Non-pollinator” Classification. (Top) Trained and tested with only directly captured images through HQ cameras, despite an imbalance in the number of pollinator and non-pollinator data points. (Bottom) Trained and tested with an increased number of non-pollinator data points obtained from the web to balance the dataset.

| *without ladybug dataset* | Class | Images | Instances | Precision | Recall | mAP50 | mAP50-95 |
| --- | --- | --- | --- | --- | --- | --- | --- |
| Validation | All | 27 | 27 | 0.934 | 0.832 | 0.871 | 0.481 |
|  | Non-pollinator | 5 | 5 | 0.966 | 0.800 | 0.800 | 0.339 |
|  | Pollinator | 22 | 22 | 0.902 | 0.864 | 0.943 | 0.622 |
| Test | All | 27 | 27 | 0.941 | 0.832 | 0.880 | 0.490 |
|  | Non-pollinator | 5 | 5 | 0.970 | 0.800 | 0.800 | 0.356 |
|  | Pollinator | 22 | 22 | 0.912 | 0.864 | 0.960 | 0.624 |
| *with ladybug dataset* | Class | Images | Instances | Precision | Recall | mAP50 | mAP50-95 |
| Validation | All | 37 | 37 | 0.841 | 0.869 | 0.923 | 0.557 |
|  | Non-pollinator | 15 | 15 | 0.861 | 0.829 | 0.888 | 0.473 |
|  | Pollinator | 22 | 22 | 0.821 | 0.909 | 0.959 | 0.641 |
| Test | All | 37 | 37 | 0.841 | 0.862 | 0.914 | 0.551 |
|  | Non-pollinator | 15 | 15 | 0.859 | 0.815 | 0.869 | 0.464 |
|  | Pollinator | 22 | 22 | 0.823 | 0.909 | 0.959 | 0.638 |

With web-sourced ladybug images to balance the volume of non-pollinator data points with pollinator data points, the YOLOv8 algorithm, trained with pollinator (bumblebee, honeybee, hoverfly, butterfly) and non-pollinator (ant, fly, ladybug) classes, showed 0.841 in Precision, 0.862 in Recall, 0.914 in mAP50, and 0.551 in mAP50-95 on the test dataset. For both pollinators and non-pollinators, precision decreased but recall increased compared to the dataset without ladybug images. Overall classification performance remained high, and non-pollinator’s mAP scores increased with the addition of ladybug images which means they can be more accurately detected from backgrounds.

Whether ladybug images were included to balance the dataset or not, both results showed classification and detection performance exceeding 80, indicating the grouping approach’s high reliability.

### 3.2 Classification of “Bee (bumblebee and honeybee),” “Hoverfly,” “Butterfly,” and “Ant”

The YOLOv8 algorithm trained with the “Bee (bumblebee and honeybee),” “Hoverfly,” “Butterfly,” and “Ant” classes showed 0.848 in Precision, 0.695 in Recall, 0.837 in mAP50, and 0.548 in mAP50-95 on the test dataset. Compared to the “Pollinator” versus “Non-pollinator” model, Precision, Recall, and mAP50 decreased, but mAP50-95 improved by 0.067 (about 7%). This indicates that while more errors were added in categorizing into the appropriate categories, the precision of object detection improved. Individually, the “Bee” class had a precision of 0.875, a recall of 1.000, an mAP50 of 0.995, and an mAP50-95 of 0.797, indicating overall good performance. However, the recall scores of “Butterfly” and “Hoverfly” were unreliable (below 0.60).

**Table 2.**
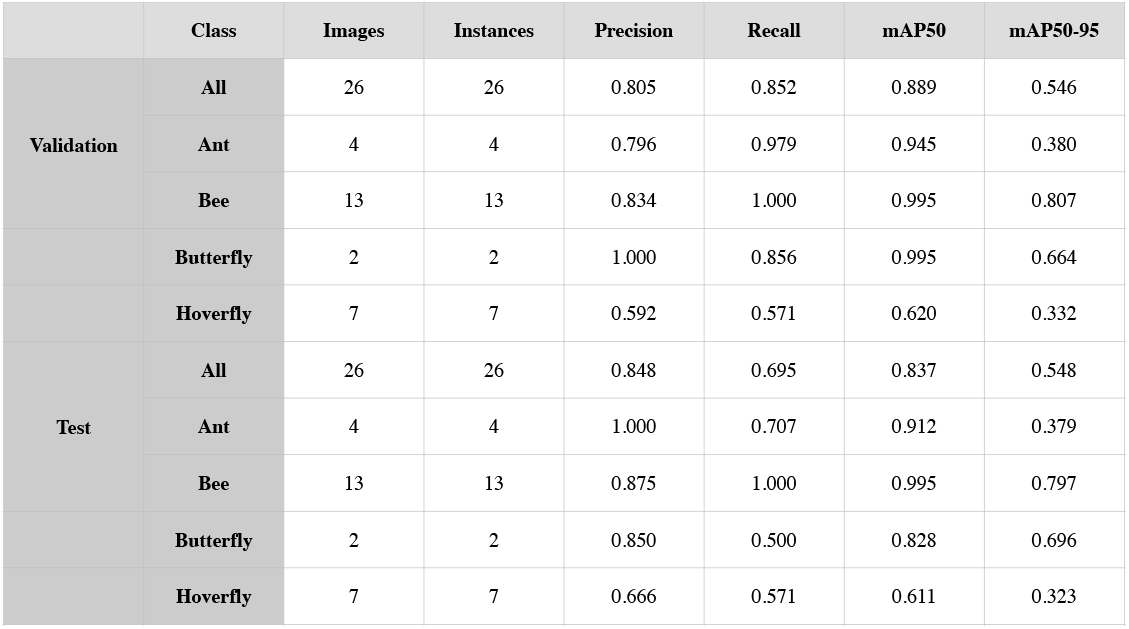
Validation and Test Classification Performance of Combining Honeybee and Bumblebee into a Single “Bee” Class, Along with Butterfly, Hoverfly, and Ant.

|  | Class | Images | Instances | Precision | Recall | mAP50 | mAP50-95 |
| --- | --- | --- | --- | --- | --- | --- | --- |
| Validation | All | 26 | 26 | 0.805 | 0.852 | 0.889 | 0.546 |
|  | Ant | 4 | 4 | 0.796 | 0.979 | 0.945 | 0.380 |
|  | Bee | 13 | 13 | 0.834 | 1.000 | 0.995 | 0.807 |
|  | Butterfly | 2 | 2 | 1.000 | 0.856 | 0.995 | 0.664 |
|  | Hoverfly | 7 | 7 | 0.592 | 0.571 | 0.620 | 0.332 |
| Test | All | 26 | 26 | 0.848 | 0.695 | 0.837 | 0.548 |
|  | Ant | 4 | 4 | 1.000 | 0.707 | 0.912 | 0.379 |
|  | Bee | 13 | 13 | 0.875 | 1.000 | 0.995 | 0.797 |
|  | Butterfly | 2 | 2 | 0.850 | 0.500 | 0.828 | 0.696 |
|  | Hoverfly | 7 | 7 | 0.666 | 0.571 | 0.611 | 0.323 |

The precision for all categories decreased by more than 10%, recall by more than 14%, and mAP50 by more than 4% compared to the Pollinator/Non-Pollinator grouping. Additionally, generally, a recall value below 0.6 for two classes is considered unsuitable for reliable use of the results. This could be because these two categories of species were rarely recorded in videos, resulting in an insufficient volume of data for effective training. This was a common occurrence across multiple solar power plants and botanic gardens, and their underrepresentation in the dataset may be typical.

Precision was highest in the order of ant (1.000), bee (0.875), butterfly (0.850), and hoverfly (0.666). Recall was highest in the order of bee (1.000), ant (0.707), hoverfly (0.571), and butterfly (0.500). The mAP50 was highest in the order of bee (0.995), butterfly (0.828), ant (0.912), and hoverfly (0.611). The mAP50-95 was highest in the order of bee (0.797), butterfly (0.696), ant (0.379), and hoverfly (0.323).

### 3.3 Classification of “Bumblebee,” “Honeybee,”“Hoverfly,”“Butterfly,” and “Ant”

The YOLOv8 algorithm trained with “Bombus,” “Honeybee,” “Butterfly,” “Hoverfly,” and “Ant” classes showed 0.856 in Precision, 0.793 in Recall, 0.864 in mAP50, and 0.554 in mAP50-95 on the test dataset. Individually, the “Bombus” class had a precision of 0.842, a recall of 1.000, an mAP50 of 0.995, and an mAP50-95 of 0.763, indicating overall good performance. However, the recall scores of “Butterfly” were unreliable (below 0.60). Precision was highest in the order of Honeybee (0.974), Ant (0.939), Bombus (0.842), Butterfly (0.792), and Hoverfly (0.732). Recall was highest in the order of Bombus (1.000), Honeybee (1.000), Ant (0.750), Hoverfly (0.714), and Butterfly (0.500). The mAP50 was highest in the order of Bombus (0.995), Honeybee (0.995), Ant (0.945), Hoverfly (0.823), and Butterfly (0.562). The mAP50-95 was highest in the order of Honeybee (0.773), Bombus (0.763), Butterfly (0.462), Ant (0.375), and Hoverfly (0.397).

**Table 3.** Validation and Test Classification Performance of Separating Honeybee and Bumblebee into Separate Classes, Along with Butterfly, Hoverfly, and Ant.

|  | Class | Images | Instances | Precision | Recall | mAP50 | mAP50-95 |
| --- | --- | --- | --- | --- | --- | --- | --- |
| Validation | All | 26 | 26 | 0.885 | 0.808 | 0.862 | 0.566 |
|  | Ant | 4 | 4 | 0.952 | 0.750 | 0.945 | 0.388 |
|  | Bombus | 6 | 6 | 0.835 | 1.000 | 0.995 | 0.793 |
|  | Butterfly | 2 | 2 | 0.822 | 0.500 | 0.538 | 0.458 |
|  | Honeybee | 7 | 7 | 0.969 | 1.000 | 0.995 | 0.773 |
|  | Hoverfly | 7 | 7 | 0.846 | 0.790 | 0.838 | 0.420 |
| Test | All | 26 | 26 | 0.856 | 0.793 | 0.864 | 0.554 |
|  | Ant | 4 | 4 | 0.939 | 0.750 | 0.945 | 0.375 |
|  | Bombus | 6 | 6 | 0.842 | 1.000 | 0.995 | 0.763 |
|  | Butterfly | 2 | 2 | 0.792 | 0.500 | 0.562 | 0.462 |
|  | Honeybee | 7 | 7 | 0.974 | 1.000 | 0.995 | 0.773 |
|  | Hoverfly | 7 | 7 | 0.732 | 0.714 | 0.823 | 0.397 |

### 3.4 Comparison Among Three Grouping Methods

The performance of three different YOLOv8 training methodologies—”Pollinator/Non-pollinator,” “Combining bumblebee and honeybee,” and “Separating bumblebee and honeybee”—was compared. Across all metrics—Precision, Recall, mAP50, and mAP50-95—the “Pollinator/Non-pollinator” model generally exhibited higher values for the overall “All” class. This indicates that the “Pollinator/Non-pollinator” grouping method is more reliable for categorizing groups and superior at detecting the presence of the target objects compared to other approaches. The “Combining bumblebee and honeybee” model and “Separating bumblebee and honeybee” model showed similar performance, but the latter generally achieved higher metrics.

Previous studies aimed at detecting and identifying pollinating insects from field recordings have focused on identifying precise taxonomic groups, primarily at the species level (Barlow *et al*., 2016; Steen, 2016; Barlow *et al*., 2017; Pegoraro *et al*., 2020; Howard *et al*., 2021; Ratanayake *et al*., 2022; Bjerge *et al*., 2022; Bjerge *et al*., 2023; Bjerge *et al*., 2024) with larger datasets containing 2,000 images (Ratnayake *et al*.), 29,960 images (Bjerge *et al*.), or 65,841 images (Hansen *et al*.). However, the results of this study suggest that a coarser grouping method proves more effective for producing an accurate classification model when the dataset images are neither fully balanced across classes nor extensive, comprising only a few thousand images. This insight is useful because, in nature, the frequency of visits by pollinating insects captured around flowers may vary by species, resulting in different captured images (Engel and Irwin, 2003).

Based on this study’s results, when data volume is insufficient, integrating these images for training not only improves classification accuracy and the reliability of generated data (e.g., counting the number of pollinators visiting flowers per unit time) but also enhances the accuracy of insect detection. This observation appears to align with the general trend in machine learning, where providing more training opportunities for individual classes can improve accuracy, particularly when the dataset size is considered insufficient (Shaikhina *et al*., 2015; Zhang and Ling, 2018; Vabalas *et al*., 2019; Kokol *et al*., 2022; Dou *et al*., 2023). Therefore, if training data is insufficient and imbalanced, employing a broader categorization method to reduce the disparity in the number of images between groups is advantageous to achieve reliable auto-monitoring and classification accuracy.

**Figure 4.**
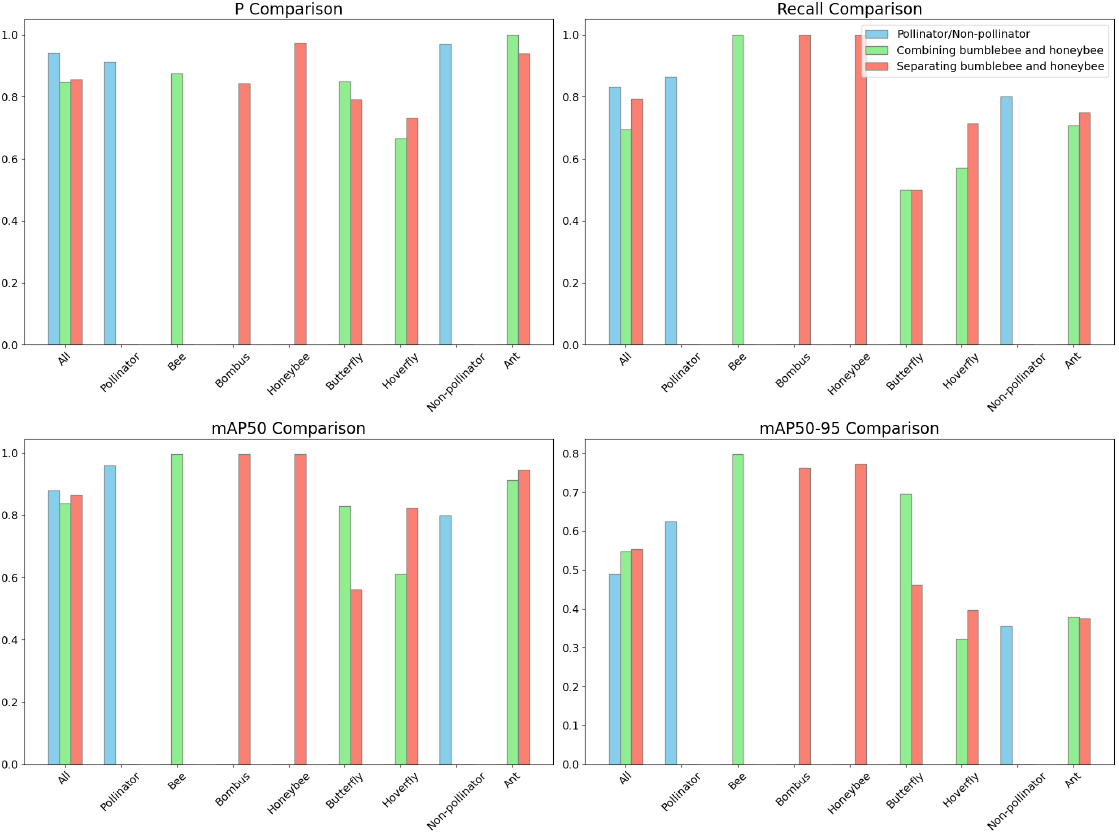
Comparison of Classification Accuracy (Precision and Recall) and Object Detection Accuracy (mAP50 and mAP50-95) Among Three Grouping Methods.

## 4. Conclusion

This study compared the performance of three YOLOv8 training methodologies: “Pollinator/Non-pollinator,” “Combining bumblebee and honeybee,” and “Separating bumblebee and honeybee.” Results showed that the “Pollinator/Non-pollinator” model generally performed better across all metrics—Precision, Recall, mAP50, and mAP50-95—indicating its superior reliability for categorizing groups and detecting target objects. The “Combining bumblebee and honeybee” and “Separating bumblebee and honeybee” models had similar performance, but the latter achieved slightly higher metrics. Previous research on detecting and identifying pollinating insects from field recordings has primarily focused on identifying precise taxonomic groups with large datasets. However, this study suggests that a coarser grouping method is more effective for accurate classification when datasets are smaller and imbalanced. This approach aligns with the general trend in machine learning, where providing more training opportunities for individual classes can improve accuracy, particularly with insufficient data. Thus, using broader categorization methods to balance the number of images between groups can enhance the reliability of automated monitoring and classification accuracy when training data is limited.

## Reference

1. Anderson, D. M. (2009). Approaches to monitoring, control and management of harmful algal blooms (HABs). Ocean & coastal management, 52(7), 342-347.

2. Barlow, S. E., Pavlik, B., Barlow, S. E., & Pavlik, B. M. (2016). Using Rana to Screen Plant Species for Effective Pollinator Support During Ecosystem Restoration. U.S.D.I. Bureau of Land Management.

3. Barlow, S. E., Wright, G. A., Ma, C., Barberis, M., Farrell, I. W., Marr, E. C., Brankin, A., Pavlik, B. M., & Stevenson, P. C. (2017). Distasteful Nectar Deters Floral Robbery. Current biology : CB, 27(16), 2552–2558.e3. 10.1016/j.cub.2017.07.012

4. Bjerge, K., Alison, J., Dyrmann, M., Frigaard, C. E., Mann, H. M., & Høye, T. T. (2023). Accurate detection and identification of insects from camera trap images with Deep Learning. PLOS Sustainability and Transformation, 2(3). 10.1371/journal.pstr.0000051

5. Bjerge, K., Frigaard, C. E., & Karstoft, H. (2023). Object Detection of Small Insects in Time-Lapse Camera Recordings. Sensors (Basel, Switzerland), 23(16), 7242. 10.3390/s23167242

6. Bjerge, K., Mann, H. M. R., Høye, T. T., Karstoft, H. (2024). A deep learning pipeline for time-lapse camera monitoring of floral environments and insect populations. 10.1101/2024.04.12.589205

7. Bjerge, K., Mann, H.M.R. and Høye, T.T. (2022), Real-time insect tracking and monitoring with computer vision and deep learning. Remote Sens Ecol Conserv, 8:315–327. 10.1002/rse2.245

8. Cayenne Engel, E., & Irwin, R. E. (2003). Linking pollinator visitation rate and pollen receipt. American journal of botany, 90(11), 1612–1618. 10.3732/ajb.90.11.1612

9. Dou, B., Zhu, Z., Merkurjev, E., Ke, L., Chen, L., Jiang, J., … & Wei, G. W. (2023). Machine learning methods for small data challenges in molecular science. Chemical Reviews, 123(13), 8736–8780

10. Howard, S. R., Nisal Ratnayake, M., Dyer, A. G., Garcia, J. E., & Dorin, A. (2021). Towards precision apiculture: Traditional and technological insect monitoring methods in strawberry and raspberry crop polytunnels tell different pollination stories. PLOS ONE, 16(5). 10.1371/journal.pone.0251572

11. Kokol, P., Kokol, M., & Zagoranski, S. (2022). Machine learning on small size samples: A synthetic knowledge synthesis. Science Progress, 105(1), 00368504211029777

12. Pegoraro, L., Hidalgo, O., Leitch, I. J., Pellicer, J., & Barlow, S. E. (2020). Automated video monitoring of insect pollinators in the field. Emerging topics in life sciences, 4(1), 87–97. 10.1042/ETLS20190074

13. Ratnayake, M. N., Amarathunga, D. C., Zaman, A., Dyer, A. G., & Dorin, A. (2022). Spatial monitoring and insect behavioural analysis using computer vision for precision pollination. International Journal of Computer Vision, 131(3), 591–606. 10.1007/s11263-022-01715-4

14. Shaikhina, T., Lowe, D., Daga, S., Briggs, D., Higgins, R., & Khovanova, N. (2015). Machine learning for predictive modelling based on small data in biomedical engineering. IFAC-PapersOnLine, 48(20), 469–474

15. Steen, R. (2017), Diel activity, frequency and visit duration of pollinators in focal plants: in situ automatic camera monitoring and data processing. Methods Ecol Evol, 8:203–213. 10.1111/2041-210X.12654

16. Vabalas, A., Gowen, E., Poliakoff, E., & Casson, A. J. (2019). Machine learning algorithm validation with a limited sample size. PloS one, 14(11), e0224365

17. Zhang, Y., & Ling, C. (2018). A strategy to apply machine learning to small datasets in materials science. Npj Computational Materials, 4(1), 25.

